# p1/s1, a 3’-nucleotidase/nuclease, allows *Leishmania major* to circumvent host innate immune response mechanisms

**DOI:** 10.1101/2025.08.15.670482

**Authors:** Stella M. Schmelzle, Michaela Bergmann, Bianca Walber, Jamal Shamsara, Tanja Ziesmann, Ute Distler, Csaba Miskey, Liam Childs, Peter Kolb, Stefan Tenzer, Katrin Bagola, Ger van Zandbergen

## Abstract

3’-nucleotidases/nucleases, distinct class I nucleases of protozoan parasites, play a pivotal role in extracellular purine salvage. As *Leishmania* are purine auxotrophs and lack *de novo* synthesis, ectoenzymes facilitating nucleotide and nucleic acid cleavage are indispensable for subsequent uptake. Employing quantitative proteomics, we identified a class I nuclease *p1/s1* cluster in *L. major* that comprises enzymes exhibiting dual 3’-nucleotidase and endonuclease activity. Expression of these enzymes is induced upon miltefosine or staurosporine treatment and was specifically detected in stationary-phase, but not in logarithmic-phase promastigotes.

After confirming secretion of p1/s1, ecto-enzymatic activity was detected on parasites and in the culture supernatant. Viable null mutants deficient for the *p1/s1* cluster were only obtained when a diCre-based inducible knockout system was applied, whereas direct deletion approaches were lethal. The viable knockout strains exhibited significantly reduced 3’-nucleotidase/nuclease activity. Notably, these parasites adapted by compensatory enrichment of various alternative purine salvage proteins at the proteomic level.

Furthermore, both enzymatic functions implied mechanisms of host-pathogen interactions to facilitate infection establishment: Utilizing 3’-nucleotidase activity, *Leishmania* generate extracellular adenosine to suppress inflammatory cytokine secretion from macrophages and reduce lymphocyte proliferation in a human primary cell model. The presence of ecto-nucleases also allowed these parasites to degrade and survive neutrophil extracellular traps, a potent first-line innate immune mechanism in pathogen defense. In summary, our integrative approach combining proteomics, immunological and genome editing methods expands current knowledge about *Leishmania major* 3’-nucleotidases/nucleases. By offering new insights into the diverse involvements in host-pathogen interactions, we highlight p1/s1 as pivotal factor during infection and potential drug target.

## Introduction

Infection of mammals with the single-cell protozoan *Leishmania* causes the neglected tropical disease leishmaniasis. *Leishmania (L.) major*, an old-world species provoking cutaneous leishmaniasis, are exposed to complex environmental changes during its diphasic life cycle. They alternate between the motile promastigote stage in the sand fly digestive tract and the intracellular amastigote stage, residing in phagolysosomes of mammalian phagocytes, mainly macrophages. Like all trypanosomatids, *Leishmania* lack purine *de novo* biosynthesis pathways, are thus auxotrophs and dependent on salvage from their host [1, 2].

In order to mobilize extracellular nucleotides and nucleic acids for metabolism, trypanosomatids utilize distinct ecto-enzymes with dual 3’-nucleotidase/nuclease activity, followed by nucleoside transport across the membrane. These class I nucleases have been biochemically characterized in a wide range of species and are exclusive for protozoans, plants, and fungi [3]. They rely on five conserved domains for Zn^2+^-mediated nucleophilic hydrolysis to cleave nucleic acids and 3’-nucleotides, with a preference for single-stranded substrates and 3’-AMP, respectively [4, 5, 3, 6].

3’-nucleotidases/nucleases are often expressed in specific life stages and can be induced by nutrient starvation in different *Leishmania* species [7–10]. For *L. major*, the amastigote-specific LmaC1N [11] and a promastigote-specific 3’-nucleotidase/nuclease [12] with membrane localization are described. p1/s1 (*LmjF.30.1500* and *LmjF.30.1510*), termed by its similarity to nucleases P1 of *Penicillium citrinum* and S1 of *Aspergillus oryzae* [5, 13], is arranged in a genomic array with four gene copies of another putative class I nuclease (*LmjF.30.1460 – 90*). The latter one is differentially transcribed in *L. major* life stages, predominantly in amastigotes [14]. Specifically, p1/s1 has been detected in the secretome of *L. major* [15].

Such nucleotidase and nuclease ecto-activities suggest a dual role in both parasite metabolism and host-parasite interactions. 3’-AMP is an intermediate of the extracellular 2’,3’-cyclic AMP pathway and is found in various tissues [16, 17]. Hence, parasites can generate extracellular adenosine by cleaving 3’-AMP through their 3’-nucleotidase activity, which triggers anti-inflammatory responses in macrophages via A_2a_ and A_2b_ receptors, thereby promoting infection establishment [9, 18]. Additionally, an endonuclease activity enables parasites to degrade extracellular DNA, facilitating escape from neutrophil extracellular traps (NETs) [19, 20]. As neutrophils act as a “Trojan horse” in early infection [21], this evasion is crucial for *Leishmania* to avoid NET-mediated immune responses.

Given the obligatory nature of purine salvage in trypanosomatids, proteins of this pathway have been declared as promising candidates for drug development. Allopurinol, a nucleobase analogue, serves as an example already used in veterinary medicine for treating canine leishmaniasis [22]. The exploitation of purine nucleoside derivatives as antiprotozoan agents supports this approach [23–25]. While the inhibition of 3’-nucleotidases/nucleases as a therapeutic strategy remains ambiguous due to the lack of specific inhibitors, the diverse roles of these enzymes at both the parasite and host levels demand further investigation.

Here, we investigated in detail the role of the ecto-3’-nucleotidase/nuclease p1/s1 in *L. major* promastigotes by employing quantitative proteomics, genome editing and *in vitro* human infection models. Besides characterizing localization and life-stage specificity of its enzymatic activity, we observed p1/s1-dependent functions to obviate innate host immune responses. Additionally, we show how *L. major* compensated an induced *p1/s1* knockout by enhancing purine salvage protein expression.

## Author summary

In this study, we describe p1/s1, a secreted 3’nucleotidase-nuclease in *Leishmania major*, to be expressed in stationary-phase but not logarithmic-phase promastigotes. Employing a quantitative proteomics dataset, we show that it is also induced upon drug-induced stress. Genomic depletion of the *p1/s1* cluster reduced both 3’-nucleotidase and endonuclease activity and impaired the parasite’s ability to survive NET-induced killing. However, *L. major^Δp1/s1^* compensate by enriching other purine salvage-linked proteins. Addition of 3’-AMP as p1/s1 substrate to a human primary macrophage infection model replicates the anti-inflammatory and lymphocyte proliferation suppression effects of adenosine. Collectively, these findings emphasize the relevance of 3’-nucleotidase/nuclease activity in driving *L. major* pathogenicity through its multiple molecular mechanisms.

## Methods

### *L. major* parasite culture

*Leishmania major* (MHOM/IL/81/FEBNI) promastigotes were cultured on rabbit blood-agar as described before [26]. Logarithmic-phase (logPh) promastigotes, 3 days post passaging, or stationary-phase (statPh) promastigotes, 6-8 days post passaging, were used for experiments. Axenic amastigotes were generated as described in [27] and washed in tissue culture plates prior to experiments. Parasites either expressed DsRed as fluorescent marker or a Cas9/T7 RNA polymerase [28]/ dimerizable Cre (diCre) construct [29]. Parasites were cultured with respective selection markers: 30 µg/ml Hygromycin B Gold, 10 µg/ml Blasticidin, 40 µg/ml G418, 40 µg/ml Puromycin (all InvivoGen), 100 µg/ml Bleomycin (Jena Bioscience).

### *L. major* proteomics

#### Sample preparation

25×10^6^ viable *L. major* promastigotes were lysed in 40 µL SDS-containing buffer (2 % (w/v) SDS, 5mM DTT, 1x cOmplete Protease Inhibitor Cocktail-EDTA (PIC, Roche), 50 mM HEPES pH 8.0). Seven biological replicates from consecutive passages were taken. To promote lysis, samples were incubated at 95°C for 5 min followed by sonication for 15 min (30 s on/off cycles, high power) at 4 °C using a Bioruptor device (Diagenode, Liège, Belgium). The protein concentration was determined using the Pierce 660 nm protein assay (Thermo Fisher Scientific) according to the manufactureŕs protocol. Subsequently, proteins (corresponding to 10 µg) were digested using single-pot solid-phase-enhanced sample preparation (SP3) as described in detail before [30, 31]. In brief, proteins were reduced and alkylated using DTT and iodoacetamide (IAA), respectively. Afterwards, 2 µL of carboxylate-modified paramagnetic beads (Sera-Mag SpeedBeads, GE Healthcare, 0.5 μg solids/μL in water as described by [30] were added to the samples. After adding acetonitrile to a final concentration of 70 % (v/v), samples were allowed to settle at room temperature for 20 min. Subsequently, beads were washed twice with 70 % (v/v) ethanol in water and once with acetonitrile. Beads were resuspended in 50 mM NH_4_HCO_3_ supplemented with trypsin (Trypsin Gold, Mass Spectrometry Grade, Promega) at an enzyme-to-protein ratio of 1:25 (w/w) and incubated overnight at 37 °C. After overnight digestion, acetonitrile was added to the samples to reach a final concentration of 95 % (v/v) followed incubation at room temperature for 20 min. To increase the yield, supernatants derived from this initial peptide-binding step were additionally subjected to the SP3 peptide purification procedure as described before [30]. Each sample was washed with acetonitrile. To recover bound peptides, paramagnetic beads from the original sample and corresponding supernatants were pooled in 2 % (v/v) dimethyl sulfoxide (DMSO) in water and sonicated for 1 min. After 2 min of centrifugation at 12,500 rpm and 4 °C, supernatants containing tryptic peptides were transferred into a new vial for MS analysis and acidified with 0.1 % (v/v) formic acid.

Alternatively, pellets were resuspended in 7 M Urea, 2 M Thiourea, 2% CHAPS subjected to tryptic digestion and processed as in [32].

#### Liquid-chromatography mass spectrometry (LC-MS)

Peptides were analyzed by LC-MS using an Evosep One chromatography system (Evosep, Odense, Denmark) coupled online to a timsTOF SCP mass spectrometer (Bruker Corporation, Billerica, MA, USA). For MS-analysis, samples were loaded onto EvoTips according to the manufactureŕs protocol. Subsequently, peptides were separated using a reversed phase C18 column (Aurora ELITE UHPLC emitter column, 15 cm x 75 µm 1.7 µm, IonOpticks) and the “Whisper Zoom 40 samples per day (40SPD)” method provided by the manufacturer. Column was heated to 50°C. Mobile phase A was 0.1% FA (v/v) in water and mobile phase B 0.1% FA (v/v) in ACN. Eluting peptides were analyzed in positive mode ESI-MS using parallel accumulation serial fragmentation (PASEF) enhanced data-independent acquisition mode (DIA) [33]. The dual TIMS (trapped ion mobility spectrometer) was operated at a fixed duty cycle close to 100% using equal accumulation and ramp times of 100 ms each spanning a mobility range from 1/K_0_ = 0.7 Vs cm^−2^ to 1.3 Vs cm^−2^. We defined 14 isolation windows with variable width covering a precursor mass range from *m/z* 307 to 1,198. Window scheme was generated using the python package py_diAID [34] resulting in seven diaPASEF scans per acquisition cycle (total cycle time of 0.85 s). The collision energy was ramped linearly as a function of the mobility from 59 eV at 1/K_0_ = 1.6 Vs cm^−2^ to 20 eV at 1/K_0_ = 0.6 Vs cm^−2^.

Alternatively, samples were measured by high-resolution nanoUPLC separation and analyzed by data-independent acquisition on the Waters Synapt G2-S platform [32].

All samples were measured in triplicates.

#### Raw data processing

MS raw data were processed using DIA-NN (version 1.9.2) [35] applying the default parameters for library-free database search. Data were searched using a custom compiled database containing UniProtKB/TrEMBL entries of the Leishmania major proteome and a list of common contaminants (version release January 2025, 8,537 entries, Taxon ID 5664). For peptide identification and in-silico library generation, trypsin was set as protease allowing one missed cleavage. Carbamidomethylation was set as fixed modification and the maximum number of variable modifications was set to zero. The peptide length ranged between 7–30 amino acids. The precursor *m/z* range was set to 300–1,800, and the product ion *m/*z range to 200–1,800. As quantification strategy we applied the “QuantUMS (high accuracy)” mode with RT-dependent median-based cross-run normalization enabled. We used the build-in algorithm of DIA-NN to automatically optimize MS2 and MS1 mass accuracies and scan window size. Peptide precursor FDRs were controlled below 1 %.

Alternatively, processing of raw data was accomplished by the ProteinLynx Global Server for *Leishmania* reference proteome and quantified using the software pipeline ISOQuant [32].

#### Data availability

The mass spectrometry proteomics data have been deposited to the ProteomeXchange Consortium (http://proteomecentral.proteomexchange.org) via the jPOST partner repository [36] with the dataset identifiers PXD066928 (ProteomeXchange) and JPST003974 (jPOST). All source code and data files are available from the authors upon request.

### Nucleotidase assay

Nucleotidase activity of 2×10^6^ intact *L. major* was determined by the release of free phosphate (P_i_) from 3’-AMP or 5’-AMP (both SigmaAldrich). After 3 hours of parasite incubation in 100 µl reaction buffer (116 mM NaCl, 5.4 mM KCl, 5.5 mM D-Glucose, 50 mM HEPES pH 7-7.6) [37] at 27 °C, parasites were separated into pellet and supernatant by centrifugation and 100 µM substrate added for 1 hour at 27 °C. Released P_i_ was quantified using a malachite green based phosphate assay kit (SigmaAldrich) according to instructions. A_620nm_ was measured with a ClarioStarPlus plate reader (BMG Labtech). A sample containing only substrate was used as blank, a control containing only parasites was always checked for P_i_ carryover. All reagents were tested for unspecific P_i_ content before use. Nucleotidase activity was expressed as µM released P_i_/2×10^6^ *L. major*/h.

### Nuclease assay

Nuclease activity of 2×10^6^ intact *L. major* was determined by visualizing enzymatic degradation of a DNA substrate on a 0.8% TAE-agarose (BioRed) gel stained with GelRed (biotium). After 3 hours of parasite incubation in 100 µl reaction buffer (30 mM sodium acetate pH 5.3, 100 mM NaCl, 2 mM ZnCl_2_) [7] at 27 °C, parasites were separated into pellet and supernatant by centrifugation and 500 µg substrate added for 1 hour at 27 °C. For endonuclease activity, single-stranded or double-stranded circular M13 phage genome (New England Biolabs) was used as substrate. For nuclease activity on NETs, respective NET-enriched supernatant was used. Reaction times were set as specified. Remaining DNA substrate was semi-quantified on gel images relative to a 500 µg undigested control using ImageJ 1.53c. A sample containing substrate and DNaseI (Invitrogen) was used for positive digestion control on every gel. Relative nuclease activity was calculated as difference to the undigested substrate.

### His_6_ enrichment and Western blot

12×10^6^ *L. major* logPh promastigotes were lysed in 20 µl Laemmli buffer (500 mM Tris/HCl pH 6,8, 38% glycerin, 10% SDS, 0.6 M DTT, 0,01% bromophenol blue) [38]. Corresponding culture supernatant was incubated with 15 µl HIS-Select nickel magnetic agarose beads (SigmaAldrich) for 30-60 min at room temperature in an overhead shaker and washed twice in PBS. Proteins were eluted in 20 µl Laemmli buffer at 65 °C for 10 min.

Protein samples were denatured for 5 min at 95 °C and separated on a 12% SDS-PAGE. After blotting, PVDF membranes were incubated with 1 µg/ml anti-αTubulin (GeneTex GTX628802) and 776 ng/ml anti-His_6_ (Cell Signaling Technology 2366) antibodies overnight at 4 °C, followed by 1 hour at room temperature with HRP-conjugated anti-mouse secondary antibody (Santa Cruz sc516102). The membrane was incubated with Immobilon Forte Western HRP substrate (Millipore) for 1 min at room temperature, signal was visualized in an ECL Chemostar (Intas).

### Human monocyte-derived macrophage (hMDM) generation

Monocytes were isolated from peripheral mononuclear cells (PBMCs) of anonymized healthy human donors as described before [39]. Buffy coats were provided by the DRK-Blutspendedienst Hessen GmbH (Frankfurt am Main). CD14^+^ monocytes were isolated by magnetic cell sorting (Miltenyi Biotec) and differentiated with 50 ng/ml M-CSF (R&D Systems) or 30 ng/ml GM-CSF (Sanofi) for 5-7 days at 37 °C, 5% CO_2_ in a humidified atmosphere. Further activation was done with 50 ng/ml IFNγ (Sigma) for 24 hours.

### Autologous peripheral blood lymphocyte (PBL) isolation

CD14^-^ PBMCs were stored at −80 °C for the time of macrophage differentiation and thawed upon usage [39].

### Isolation of polymorphonuclear cells (PMNs)

PMNs of healthy individuals were isolated from fresh blood as described in [40] or from buffy coats as previously described [26]. 12×10^6^ cells were seeded on poly-D-lysine (Gibco) coated 12-well plates. Neutrophil extracellular trap (NET)-enriched supernatants were induced and harvested as reported before [41]. DNA content was determined using a Quant-it RiboGreen assay (Invitrogen).

### Determination of hMDM infection

Differentiated macrophages were infected with *L. major* promastigotes from the statPh at a multiplicity of infection (MOI) of 10 for 3 hours at 37 °C before remaining extracellular parasites were washed off. After the specified time post infection (pi), the infection rate of viable single cells was quantified by detecting the parasite’s fluorescence marker by flow cytometry on either LSR Fortessa or Symphony A3 Cell Analyzer (both BD Bioscience). Parasite burden was determined as mean fluorescence intensity (MFI) of single, viable, infected macrophages.

Alternatively, cells were fixed in 4 % PFA (Sigma) and permeabilized using 0.5 % saponin (AppliChem PanReact), then intracellular parasites were stained with 1:10,000 anti-*L. major* rabbit serum and 1:1000 goat Anti-rabbit AlexaFluor™ 568 secondary antibody (Invitrogen A-11036).

If applicable, adenosine (SigmaAldrich) or 3’-AMP (Santa Cruz Biotechnology) were added during infection at indicated concentrations.

### Analysis of PBL proliferation in a co-culture with hMDM

Macrophages and PBLs were stained in 1 µM CellTrace Far Red (Invitrogen) for 20 min at 37 °C. 0.2×10^6^ stained macrophages were infected with *L. major* promastigotes from the statPh at MOI=20 for 3 hours at 37 °C before 1×10^6^ stained PBLs were added. Infection was carried out in U-shaped 96-well plates with 2×10^4^ hMDMs per well and 10 technical replicates per condition which were pooled before analysis. PBL proliferation was analyzed by flow cytometry 4 days pi as CellTrace_low_ fraction of viable, single PBLs. Infection rate and parasite burden of hMDM were determined as stated before.

If applicable, adenosine or 3’-AMP were added together during infection at indicated concentrations.

### Neutrophil killing assay

Isolated granulocytes were tested for activation status by CD66b and CD62L staining beforehand. 2×10^6^ PMNs in a 24-well plate were treated with 10 µg/ml Cytochalasin D (SigmaAldrich) for 20 min at 37 °C. 130 U DNaseI were added at the same time, if applicable. *L. major* statPh promastigotes were added at MOI=1 and incubated for 2 hours at 37 °C and then transferred to 27 °C for two days. Corresponding controls containing reagents and parasites but no PMN were treated the same. Motile, viable parasites were counted in a Neubauer chamber and numbers normalized to the respective control gave the percentage of viable parasites.

### ELISA

Cell free supernatants from infected samples were analyzed for soluble cytokine content using enzyme-linked immunosorbent assay (ELISA) (DuoSet, R&D Systems) following instructions. TNFα content was measured as 1:12.5 dilutions. Signals were developed using horseradish peroxidase substrate solution (BioLegend) and stopped by addition of 2 N H_2_SO_4_ (Carl Roth). Absorbance was measured at 450 nm using 570 nm as reference wavelength at a ClarioStarPlus. Total cytokine concentration was determined by 4-parameter regression of the standard row in Mars v4.00. Samples were measured in duplicates.

### Staining of surface markers

Human primary cells were stained for surface markers in PBS pH 7.2 plus 0.5 g/L BSA and incubated for 30 min at 4 °C. Antibodies were added accordingly: anti-A_2a_-AlexaFluor488, anti-A_2b_-AlexaFluor647 (both R&D Systems, 1 µg/1×10^6^ cells). Cells were also stained with 1:1000 ZombieAqua (BioLegend) or 1µM SYTOXBlue (Invitrogen) as viability marker. Expression was analyzed by flow cytometry as percentage positive cells.

### Generation of *L. major* inducible knockout strains

*L. major* expressing Cas9/T7 RNA polymerase and a dimerizable Cre (diCre) construct were used to integrate a 3’ and a 5’ flox cassette in two consecutive steps by CRISPR/Cas9-facilitated homologous recombination, adapting the method of [29]. Each cassette was amplified using 0.5 µM forward and reverse primer (sequences see Tab. 1), 15 ng/µl plasmid template (pGL2314 for 3’ flox [29], pTNeo-eGFP-loxP (modified based on [28]) for 5’ flox), 4 µl GC Enhancer, 10 µl Platinum Mastermix (both Invitrogen) and 3 µl H_2_O in a S1000 Thermal Cycler (BioRad) set to 35 cycles of 10 s at 98 °C and 3 or 4 min (3’ flox/ 5’ flox) at 72 °. Single guide RNA (sgRNA) templates were amplified using 2 µM primer, 2 µM scaffold, 5 µl PWO Mastermix (Roche) and 1 µl H_2_O for 35 cycles of 10 s at 98 °C, 30 s at 60 °C and 15 s at 72 °C. Products were checked on a 1% agarose gel and sterilized for 5 min at 94 °C.

**Tab. 1:**
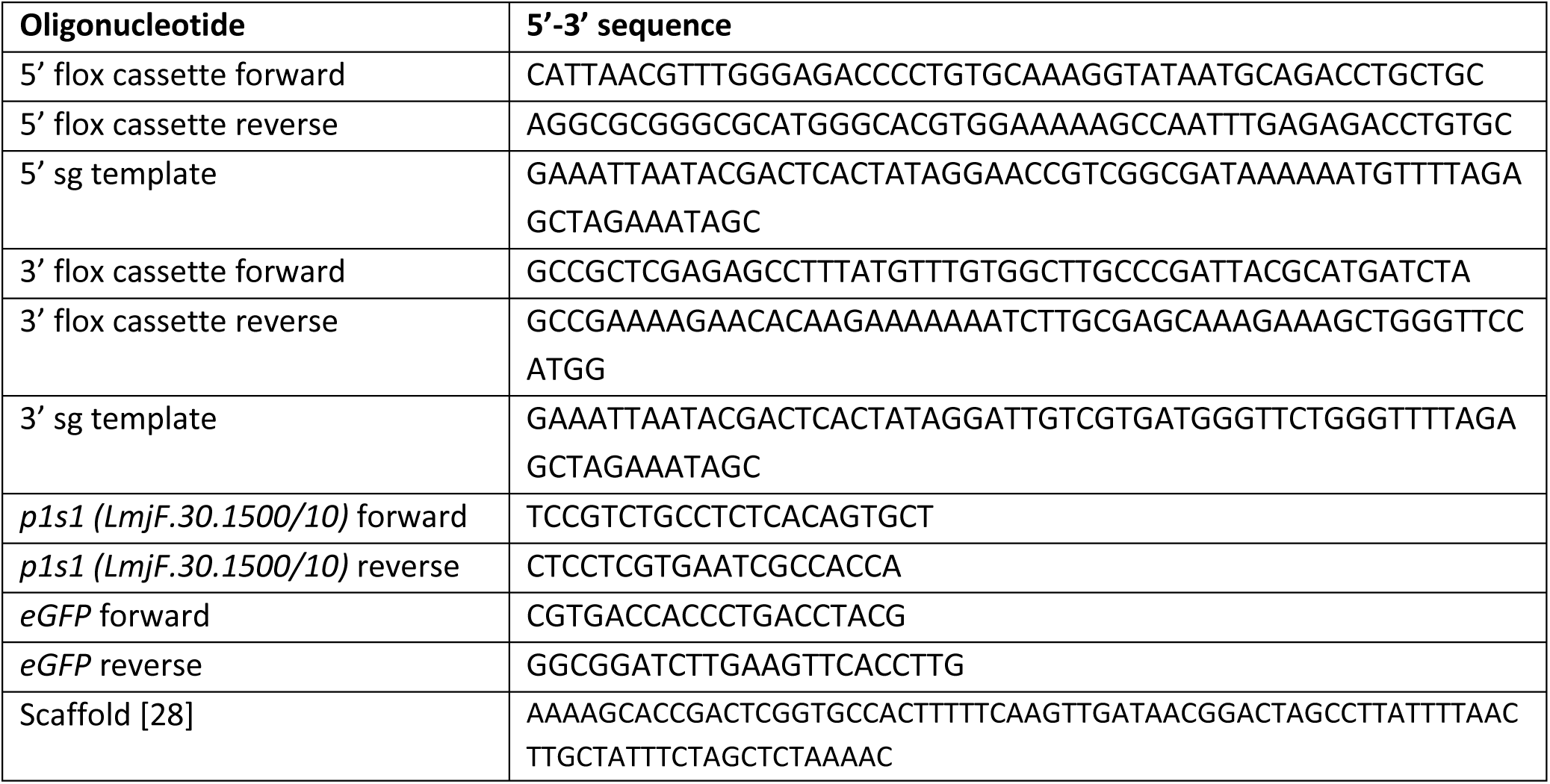
Oligonucleotide sequences.

60×10^6^ promastigotes from the logPh were pulsed once with the PCR products in Human T-cell Nucleofector® buffer (Lonza) using a Nucleofector® 2b (Lonza) with program X-001. Electroporated cells were incubated overnight in 9 ml *L. major* medium at 27 °C. After addition of the respective antibiotics and 10% fetal calf serum (SigmaAldrich), parasites were distributed in 100 µl on a 96-well rabbit blood agar plate. After 14-21 days, grown clones were checked for mCherry (3’ flox) and eGFP (5’ flox) fluorescence using flow cytometry and single cell clones were generated by serial dilution.

Recombination in full-floxed strains was induced by addition of 100 nM rapamycin (SigmaAldrich) and fluorescence monitored over 9 days. Single cell clones were generated as above and checked for *p1/s1* and *eGFP* status by PCR (2 µl genomic DNA, 1.5 µl forward and reverse 10 µM primer each, 5 µl PWO Mastermix) by 3 min at 95 °C and 25 cycles of 30 s at 95 °C, 30 s at 59 °C, 45 s at 72 °C plus final extension for 10 min at 72 °C. Amplified products were checked on a 1 % agarose gel.

### Generation of *L. major* addback strains

*p1/s1* (*LmjF.30.1500/10*) was inserted without protein tag into the multiple cloning site of pLexLm plasmid, a plasmid based on pLEXSY-ble2.1 (Jena Bioscience, Cat. No. EGE-271) in which the rRNA promotor region of *L. donovani* has been exchanged by an *L. major* rRNA promotor region. 60×10^6^ *L. major^Δp1/s1^* clone E10 logPh promastigotes were electroporated with SwaI (New England Biolabs) linearized and sterilized plasmid. Electroporated cells were incubated overnight in 9 ml *L. major* medium at 27 °C. After addition of bleomycin and 10% FCS, parasites were spread in 100 µl on a 96-well rabbit blood agar plate. After 14-21 days, grown clones were checked for *p1/s1* status using PCR and single cell clones were generated.

### Cell death assays

*L. major* logPh promastigotes were treated with increasing concentrations of miltefosine (Calbiochem) for 48 h, and parasites were used in different assays to quantify different hallmarks of cell death:

Metabolic rate of 5×10^6^ promastigotes was determined by colourimetric turnover of 1.1 mM Thiazolyl blue tetrazolium bromide (MTT, abcam) in phenolred-free RPMI (SigmaAldrich) for 4 hours at 27 °C. Formazan crystals were dissolved in DMSO (AppliChem PanReact) and absorbance was measured at 540 nm, using 700 nm as reference wavelength, at a ClarioStarPlus. Values were blanked to a sample containing parasites but no MTT. Relative cell metabolism representing viability was calculated by normalizing to the DMSO control.

2×10^6^ promastigotes were stained with 1:1000 AnnexinV-FITC (Miltenyi Biotec) in Ringer solution (B. Braun) for 20 min at 4 °C. Labelling was analyzed by flow cytometry, gating was set using a control stained in calcium-free PBS.

2×10^6^ promastigotes were permeabilized in 70 % (v/v) EtOH for 30 min at −20 °C and stoichiometrically stained for DNA content using 5 µM SYTOXGreen (Invitrogen) in PBS containing 50 mM EDTA pH 7.4 and RNase (Thermo Fisher) for 30 min at 37 °C. Parasites were analyzed by flow cytometry on LSR II SORP (BD Bioscience), gating was determined on an unstained control. The share of cell cycle phases was determined based on DNA content.

Effective concentration (EC)_50_ values were calculated based on 4-parameter regression in GraphPad Prism with constraints for bottom and EC_50_ ≥ 0 and stated as ± 95% confidence interval (CI).

### Statistics

All graphs display mean ± standard deviation and were generated in GraphPad Prism v9.5.0. Significance was determined using a two-sided Welch’s t-test with * p ≤ 0.05, ** p ≤ 0.01, *** p ≤ 0.001, **** p ≤ 0.0001.

## Results

### A set of 9 proteins is commonly enriched in *L. major* upon cell death pressure, including p1/s1

Identification of stress-related proteins in *Leishmania* is of great interest as they can feature potential drug targets [44–46] or be involved in mediating apoptotic-like cell death, a mechanism used by *Leishmania* to promote disease development and circumvent effective adaptive immune responses [39, 47]. In order to designate such proteins in *L. major*, we applied stress to the parasites by treatment with sublethal doses of miltefosine, an antileishmanial drug, for 3 hours or staurosporine, an unspecific kinase inhibitor, for 6 hours. Following treatment, we performed a quantitative proteome analysis to compare protein expression (Fig1A-B). The treatment concentrations were set to sublethal doses leading to cell death after 24 hours (SuppFig. 1). Additionally, stationary-phase (statPh) promastigotes, harboring parasites that undergo intrinsic cell death, were analyzed and compared to untreated logPh parasites (Fig. 1C). By choosing these conditions, proteins of immediate stress responses prior to cell death or survival promotion can be detected in the intersection of enriched hits. Proteins involved in drug-specific responses or metacyclogenesis of the statPh are solely detected in single conditions and are thus excluded.

**Figure 1.**
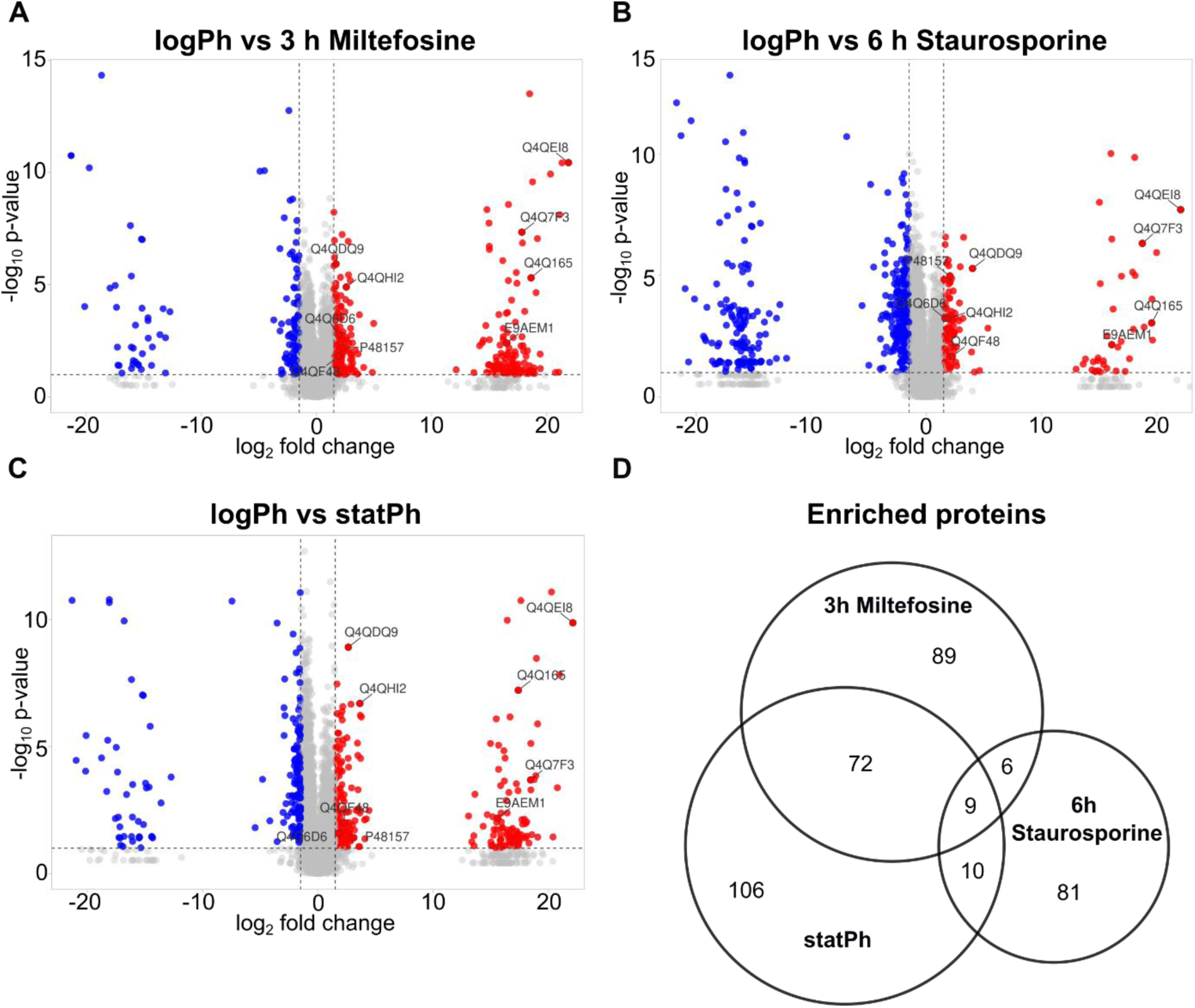
Quantitative proteomic analysis of cell-death induced *L. major*. **A**- Volcano plot of quantified proteome of *L. major* treated with 25µM Miltefosine for 3 h, **B**- 25µM Staurosporine for 6 h and **C**- statPh promastigotes compared to untreated, logPh promastigotes. Volcano plot visualized with VulcaNoseR [42], commonly enriched proteins are labelled (Tab. 2). Cutoffs were set at log_2_FC <-1.5, >1.5 and p-value < 0.01. **D**- Venn diagram of protein numbers significantly enriched in the different comparisons. Template created using [43]. Analysis based on 7 biological replicates and 3 technical replicates each.

**Tab. 2:**
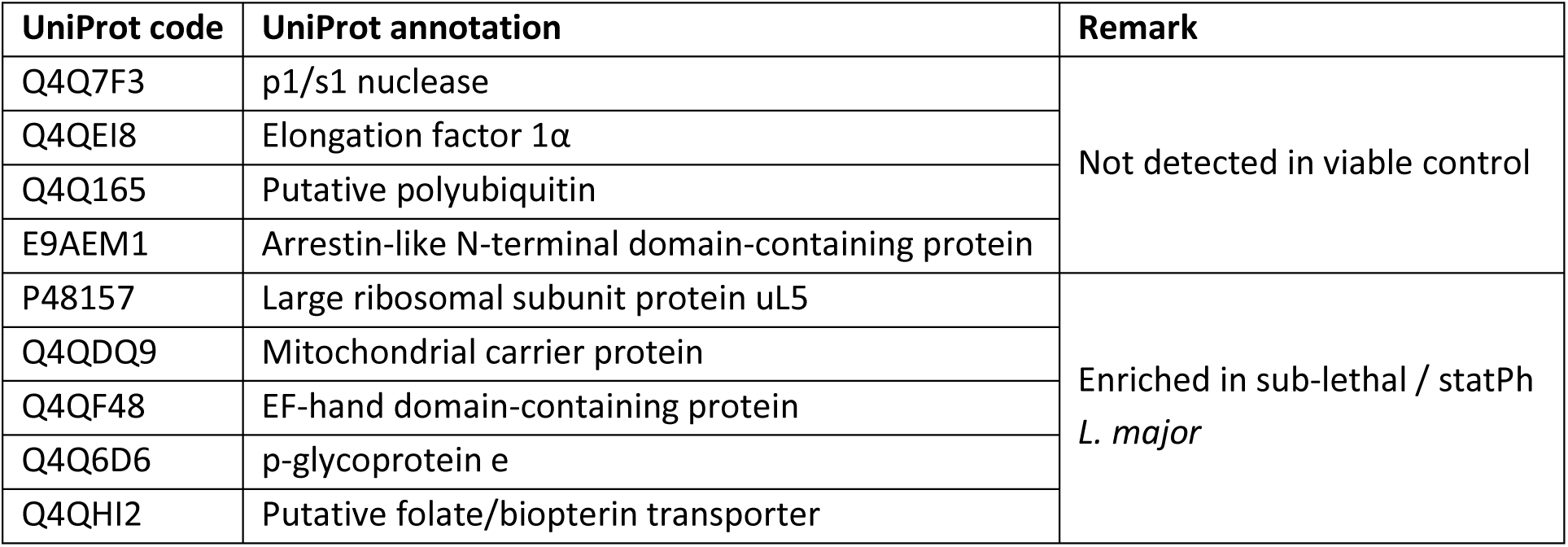
Proteins commonly enriched in *L. major* undergoing cell death; annotations extracted from UniProt [49].

Across the three comparisons, 4116 proteins were detected in total, corresponding to 51.73% of the reference *Leishmania* proteome. Taking the intersection of enriched hits, only five proteins were commonly upregulated in all three comparisons, four were exclusively detected in sub-lethal parasites or statPh promastigotes (Fig. 1D, Tab. 2). Among these hits are several proteins involved in protein biogenesis and turnover but also proteins of diverse predicted functions. One of these hits, p1/s1, is predicted as class I nuclease. In *Leishmania*, such enzymes are involved in essential purine salvage by eliciting 3’-nucleotidase and endonuclease activity and are presumably involved in cellular functions as well as in host-parasite interactions (reviewed in [48]). Given p1/s1 expression in stress-induced and infectious statPh *L. major*, this hit stands out as central link between parasite metabolism and disease progression. Studying its multiple functions and its potential role in parasite cell death could establish p1/s1 as potent drug target.

### Ecto-nucleotidase and -endonuclease activity of *L. major* increase from the logarithmic to stationary growth phase

Indeed, detected peptides assigned to p1/s1 can derive from six open reading frames arranged in a tandem array (*LmjF.30.1460* to *LmjF.30.1510)*. Sequence details were retrieved from EnsemblProtists release 61. All open reading frames share a common C-terminal domain. Whereas the four genes *1460-90* contain a putative transmembrane domain (TMD), *1500* and *1510* code for an N-terminal signal sequence for secretion (Fig. 2A), as predicted by DeepLoc 2.1 [50]. We verified secretion by overexpressing a C-terminally His_6_-tagged *LmjF.30.1500* construct in *L. major*^Cas9/T7/eGFP^. Parasite lysate was used for SDS-PAGE, whereas the corresponding culture supernatant was used for His_6_ enrichment over magnetic Ni-NTA beads. This eluate was also separated by SDS-PAGE. p1s1-His_6_ and the housekeeping protein α-Tubulin were detected by Western blot. Whereas no His_6_-tagged proteins were detected in the lysate, a His_6_-tagged protein approximately at the predicted size of p1/s1 (35.1 kDa including signal sequence) was visible in the culture supernatant for both overexpressing clones H9 and F3 (Fig. 2B), confirming p1/s1 secretion.

**Figure 2.**
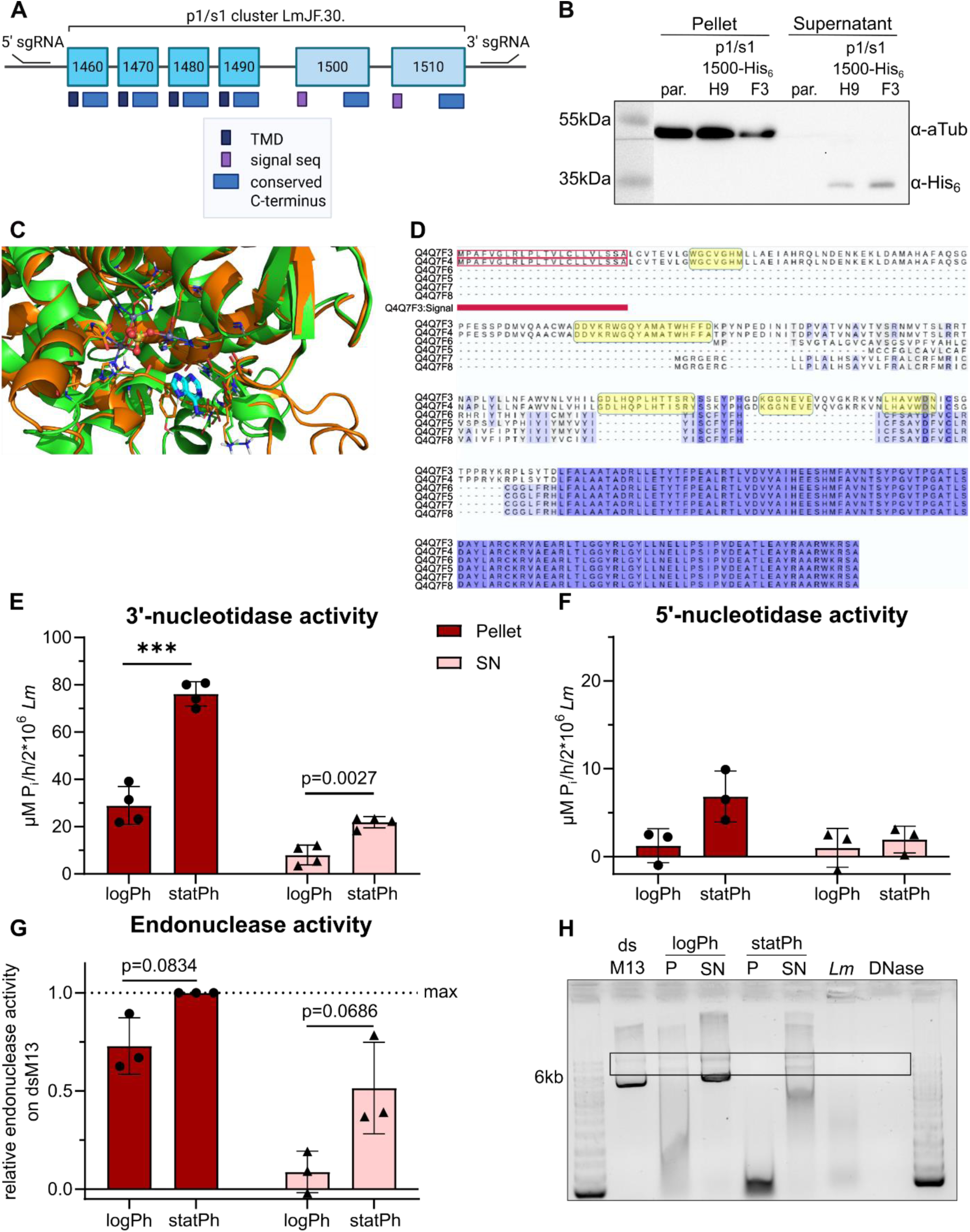
Ecto-nucleotidase and -endonuclease activities of *L. major* increase in the stationary-phase. **A**- Depiction of the genomic *p1/s1* cluster of six genes (*LmjF.30.1460 – 1510*) sharing a conserved domain but encoding either a signal sequence or a transmembrane domain (TMD). Indicated 5’sgRNA and 3’sgRNA were used for CRISPR Cas9-mediated inducible knockout of the cluster. Figure created with Biorender.com. **B**- Western blot to determine p1/s1-His_6_ localization in *L. major* promastigotes expressing a C-terminally His_6_-tagged LmjF.30.1500 construct (clones H9 and F3). Secreted, tagged proteins were enriched with magnetic Ni-NTA beads. Representative blot of n=3. **C**- Superimposition of the AlphaFold-predicted structure of Q4Q7F3 (orange, encoded by *LmjF.30.1510*) onto the experimental structure of the homologous enzyme (green; PDB ID: 3W52). The active site residues and the bound adenine base (cyan) in 3W52 are shown as sticks. Zinc ions are represented as gray spheres, and the sulfate ion trapped among the zinc ions is shown in ball-and-stick representation. **D**- Multiple sequence alignment of Q4Q7F8-F3, as encoded in the *LmjF.30.1460-1510* cluster. The regions corresponding to homologous active sites in nucleoside enzymes are highlighted in yellow boxes. **E**- 3’-nucleotidase activity of intact *L. major* was measured as P_i_ release from 3’-AMP. n=4. **F**- 5‘-nucleotidase activity of intact *L. major* was measured as P_i_ release from 5’-AMP. n=3. **G**- Endonuclease activity of 2×10^6^ total *L. major* was measured by degradation of double-stranded M13 phage genome (dsM13) for 60 min. **H**- Representative gel of n=3, box denotes area of semi-quantification.

To identify the putative homologous active site of p1/s1, AlphaFold-predicted structures of proteins coded in the *LmjF.30.1460* -*1510* cluster were superimposed on homologous template structures, neglecting the putative signal sequence (Fig. 2C). Based on homology to Endonuclease 2 from *Arabidopsis thaliana* (3W52), five regions were identified to participate in the p1/s1 active site (Fig. 2D), coinciding with previous reports on the *L. donovani* 3’-nucleotidase/nuclease homologue *Ld*Nuc^S^ [8]. In contrast to the putative membrane-bound proteins Q4Q7F8-F5, encoded by *LmjF.30.1460-90*, only the secreted enzymes Q4Q7F4-3, encoded by *LmjF.30.1500-10*, contain this active site. Beyond p1/s1, further enzymes (Q66VY6, Q4QGQ3, Q4Q630) possess a homologous active site, indicating a putative functional redundancy.

Further, to assess overall 3’-nucleotidase activity arising from ecto-enzymes of *L. major,* and to compare p1/s1 expression and 3’-nucleotidase activity patterns, we quantified the release of free phosphate (P_i_) from 3’-AMP. Intact parasites, harboring all membrane-bound ecto-enzymes, were separated from the supernatant, which contains secreted enzymes. Besides the secreted p1/s1, the *L. major* genome codes for additional 3’-nucleotidases/nucleases predicted to be membrane-localized. Comparing logPh and statPh promastigotes, a significant increase in 3’-nucleotidase activity in both the pellet and the supernatant fraction was observed, well corresponding to the elevated p1/s1 protein levels in statPh (Fig. 1C). Overall, activity associated with pelleted parasites was higher (Fig. 2E), likely related to non-secreted 3’-nucleotidases/nucleases. In contrast, degradation of 5’-AMP through 5’-nucleotidases was generally much lower than 3’-nucleotidase activity (Fig. 2F).

Accordingly, ecto-endonuclease activity was determined using double-stranded M13 phage genome (dsM13) as substrate. Its degradation was visualized on an agarose gel for semi-quantification. Similar to 3’-nucleotidase activity, endonuclease activity is increased in statPh and a higher activity was observed on pelleted parasites than in the supernatant (Fig. 2G-H). These results align with the pattern seen for 3’-nucleotidase activity and p1/s1 protein expression, suggesting both functions arise from p1/s1.

### Presence of 3’-AMP during *L. major* hMDM *in vitro* infection mimics anti-inflammatory effects of extracellular adenosine

As shown in Fig. 2C, *L. major* statPh promastigotes can hydrolyze extracellular 3’-AMP to adenosine and P_i_. Extracellular adenosine is a well-characterized anti-inflammatory stimulant of immune cells. GM-CSF-differentiated macrophages stimulated with IFNγ exhibit the highest share of cells among tested macrophage phenotypes positive for its cell surface receptors A_2a_ and A_2b_ (SupFig. 2A-B). Activation of these receptors can result in anti-inflammatory immune modulation. Thus, we examined whether macrophage exposure to adenosine or 3’-AMP during *L. major* infection can the pro-inflammatory cytokine response. Indeed, TNFα release from macrophages 3 hours post *Leishmania* infection decreases dose-dependently with increasing concentrations of adenosine. At 100 µM adenosine, TNFα levels align with those of uninfected controls (Fig. 3A and SupFig. 2D). Addition of 3’-AMP to the infection partially mimics the effect observed with adenosine.

**Figure 3.**
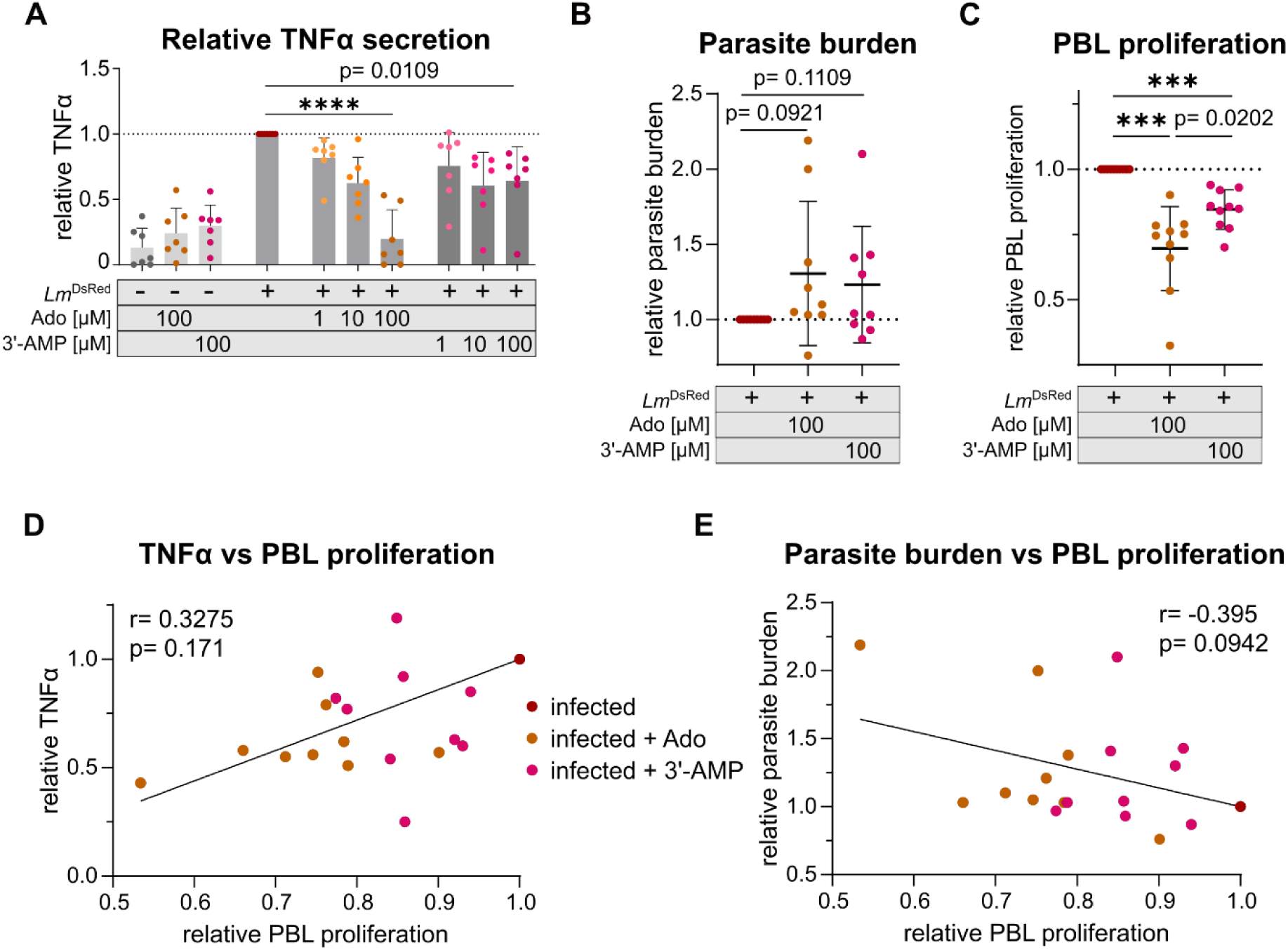
Presence of adenosine or 3‘-AMP during *L. major*^DsRed^ infection of hMDM decreases TNFα secretion and PBL proliferation. **A**- Soluble TNFα secreted from hMDM(GM-CSF+IFNγ) was determined in supernatant collected 3 hours pi using ELISA and normalized to the infected sample without additives. **B**- Relative parasite burden measured as DsRed MFI of infected hMDM(GM-CSF+IFNγ) normalised to the infected control without additives. Macrophages were infected for 3 hours with a MOI=20 of statPh *L. major*^DsRed^ in the presence of adenosine or 3‘-AMP before autologous PBLs were added to the infection and incubated for 4 days. **C**- Percentage of proliferated PBLs was determined as CellTrace_low_ lymphocytes 4 days pi and normalized to the infected control without additives. **D**- Pearson correlation of relative PBL proliferation and TNFα secretion and **E**- relative PBL proliferation and parasite burden. A: n=7 donors in 4 independent experiments; B-E: n=9-10 donors in 4 independent experiments.

TNFα is a known driver of the antileishmanial host response [51]. Neutralization of TNFα using specific monoclonal antibodies was previously shown to reduce *Leishmania*-induced proliferation of autologous peripheral blood lymphocytes (PBLs) in a hMDM:PBL coculture [52]. T cells are the predominant cell type within PBLs. Thus, we tested whether extracellular adenosine exposure impacts CD14^-^ PBL proliferation 4 days post infection. Although parasite burden of macrophages was not significantly affected, an underlying trend towards an increased parasite burden upon addition of adenosine or 3’-AMP was observed (Fig. 3B). Relative to the control, reduction in proliferation of PBLs and TNFα secretion indeed was apparent for samples treated with adenosine, 3’-AMP again mimicked this (Fig. 3C). Addition of adenosine or 3’-AMP to uninfected cocultures did not affect the share of proliferating PBLs. The proportion of proliferating cells increased greatly upon infection (SupFig. 2E). Additionally, a mild to moderate correlation between relative PBL proliferation and TNFα secretion or parasite burden in infected samples was identified (Fig. 3D+E), suggesting accessory factors induced by adenosine signaling also impact the observed effects. Thus, by generating adenosine through their 3’-nucleotidase activity, *L. major* can suppress an effective early immune response.

### p1/s1 is dispensable for *L. major* but knockout mutants show reduced 3’-nucleotidase/nuclease activity

To confirm that the observed ecto-3’-nucleotidase/nuclease activity of *L. major* arises from p1/s1, the generation of knockout parasites lacking the *p1/s1* cluster was attempted. However, CRISPR-Cas9-based double homologous replacement approaches [28] failed to yield viable null mutants when sgRNAs binding downstream of *LmjF.30.1460* and upstream of *LmjF.30.1510* were used (Fig. 2A). Targeting single genes in the *p1/s1* cluster was not possible due to very high homology between coding and intergenic regions within the cluster. Neither facilitated approaches, using stable genetic integration of a *LmjF.30.1500* construct into the *SSU* locus, or nutritional complementation by supplementing 100µM adenosine to the medium, yielded viable clones (Tab. 3).

**Tab. 3:** Targeted gene deletion approaches deployed to knockout the p1/s1 cluster using sgRNA binding binding downstream of *LmjF.30.1460* and upstream of *LmjF.30.1510*.

| Deletion approach | Result |
| --- | --- |
| Gene deletion | Only heterozygous strains or lethal |
| Facilitated p1/s1 deletion with genetic complementation | Only heterozygous strains or lethal |
| Facilitated p1/s1 deletion with nutritional complementation | Only heterozygous strains or lethal |
| Inducible p1/s1 deletion | Viable null mutants |

Thus, an inducible diCre approach [29] was applied. Excision of *LmjF.30.1460-1510* was induced by addition of rapamycin, that catalyzes dimerization of the Cre recombinase in two clones (E10 and F4) carrying a floxed *p1s1* cluster (Fig. 4A). The eGFP gene of the 5’ flox casette is excised together with the gene cluster during the recombination event. Therefore, loss of eGFP signal over time was used to track successful recombination. Up to 7 days post rapamycin induction, eGFP positivity was lost continuously in both clones, while a 3’ integrated mCherry signal was retained (Fig. 4B+C). Following serial dilution to yield single-cell clones, genomic loss of *LmjF.1500/10* and *eGFP* was confirmed by diagnostic PCR (Fig. 4D+E). One confirmed *L. major^Δ^*^p1/s1^ single-cell clone per floxed strain was chosen for further experiments as induced *p1/s1* knockouts (iKOs). Based on *L. major^Δ^*^p1/s1^ iKO E10, an addback strain was established by genomic integration of *LmjF.1500* into the *SSU* locus. In comparable culture conditions, both *p1/s1* iKO strains show greatly reduced parasite density in the logPh and statPh compared to the parental strain, whereas the addback strain showed intermediate growth behavior (Fig. 4F).

**Figure 4.**
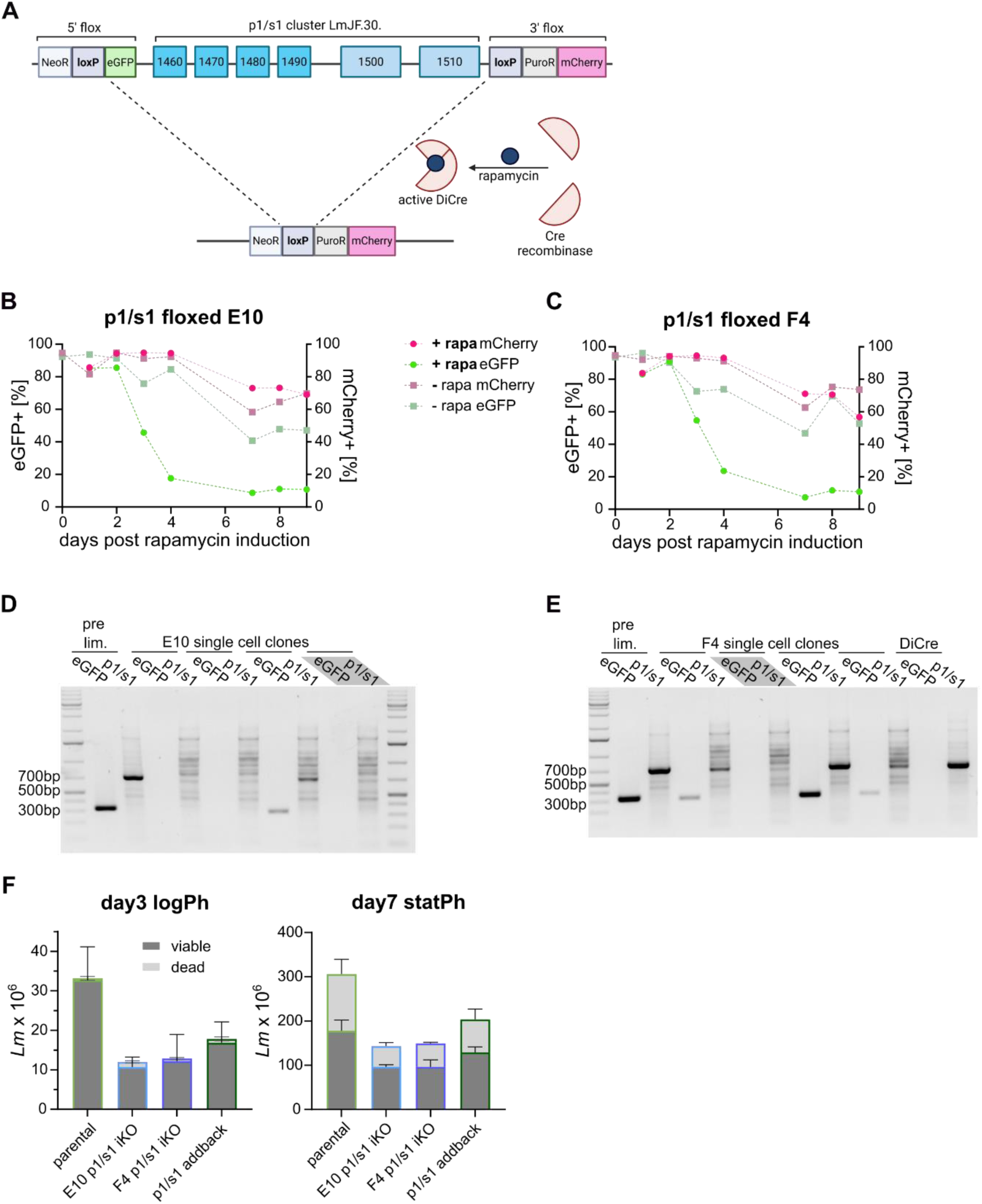
diCre based inducible knockout approach yields *L. major p1/s1* null mutants. **A**- Schematic visualization of the inducible, dimerizable Cre (diCre) system used to generate *L. major^Δ^*^p1/s1^. Full floxed parasites (top) expressing a diCre construct were induced with rapamycin to initiate Cre-mediated excision of the *p1/s1* cluster alongside the *eGFP* gene. Successful recombination (bottom) can be traced by the loss of eGFP signal. Figure created with biorender.com. **B**, **C**- eGFP and mCherry signal upon diCre induction with rapamycin measured until 9 days post induction in two p1/s1 full floxed clones E10 and F4, respectively. **D**- Diagnostic PCR to detect eGFP and p1/s1 genes in genomic DNA of single cell clones derived from clones E10 and **E-** F4 post rapamycin induction. *L. major^Δ^*^p1/s1^ clones used for further experiments are marked in gray and further named as E10 and F4 induced knockouts (iKO). *L. major*^Cas9/T7/diCre^ and induced parasites before limiting dilution (pre.dil.) were used as controls. **F**- Growth of *L. major*^Cas9/T7/diCre/3‘flox^, *L. major^Δ^*^p1/s1^ clones E10 & F4 and *L. major*^p1/s1 addback^ were counted on day 3 (logPh) and 7 (statPh) post passaging for viable and dead parasites. All strains were grown under selection pressure of the same antibiotics. n=3 of consecutive passages.

When using *L. major^Δ^*^p1/s1^ for 3’-nucleotidase activity assays, both null mutants show considerably diminished activity in the supernatant fraction, whereas the addback partly restored activity. 3’-nucleotidase activities on the pellet fraction are inconclusive but suggest no difference between parental and knockout strains (Fig. 5A). Similarly, *L. major^Δ^*^p1/s1^ exhibit declined endonuclease activity on double-stranded substrate (Fig. 5B-C) and delayed degradation of single-stranded substrate (Fig. 6D-F), both in the pellet and the supernatant fractions. *L. major^Δ^*^p1/s1^ addback promastigotes here behaved like the parental strain. Thus, behavior of the parental strain coincided with 3’-nucleotidase/nuclease activity observed in Fig. 2, whereas null mutants exhibit a distinct phenotype with delayed or reduced degradation of substrates. The *L. major* genome codes for further 3’-nucleotidases/nucleases besides p1/s1. These redundant enzymes Q66VY6, Q4Q630 and Q4QGQ3 all possess an active site homologous to p1/s1, so residual enzymatic activity in mull mutants was expected.

**Figure 5.**
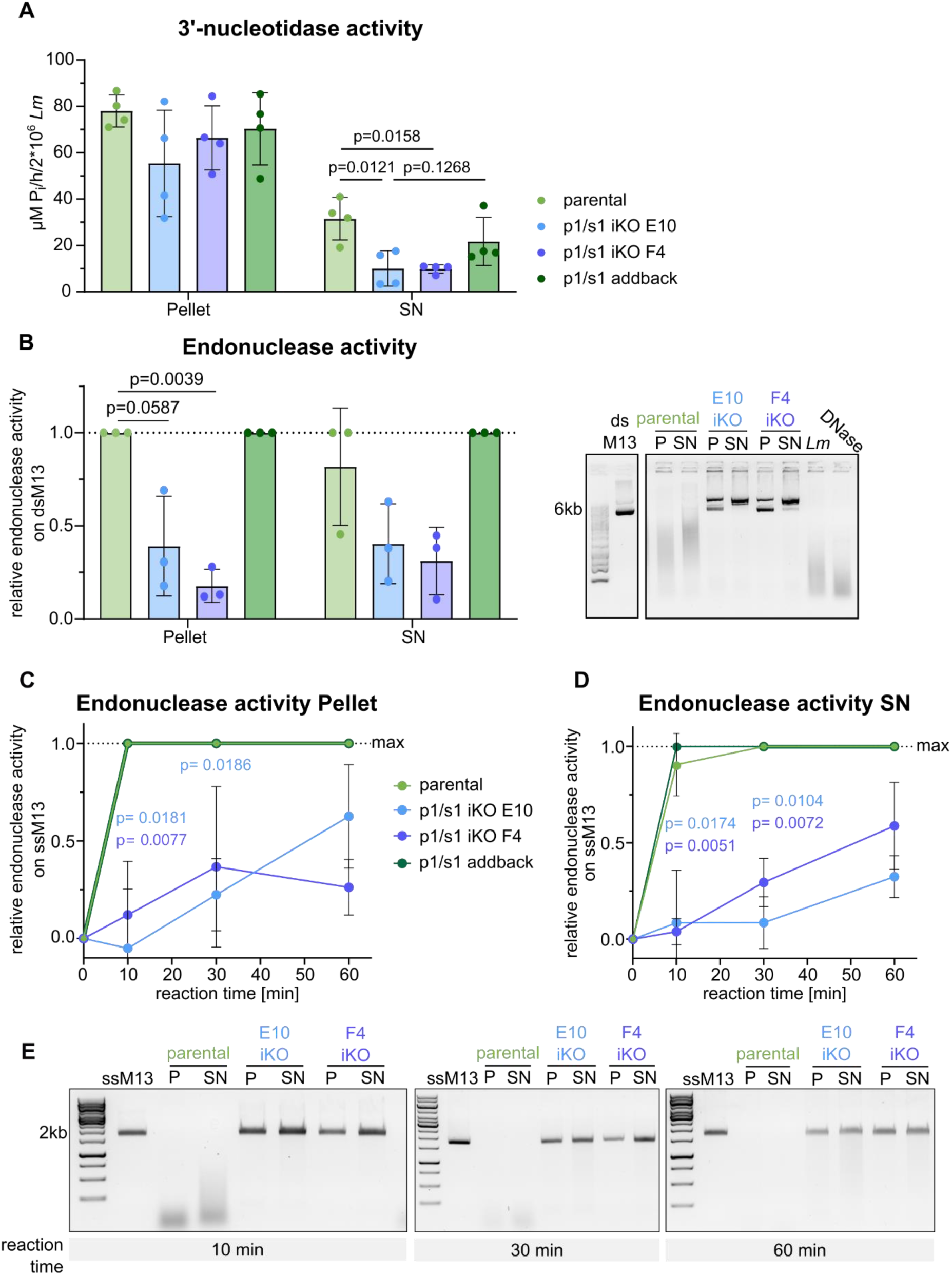
*L. major p1/s1* null mutants show reduced 3’-nucleotidase/nuclease activity. **A**- 3‘-nucleotidase activity was measured by the release of P_i_ from 3‘-AMP on 2×10^6^ total *L. major* statPh promastigotes (*L. major*^Cas9/T7/diCre/3‘flox^ *L. major^Δ^*^p1/s1^ clones E10 & F4 and *L. major*^p1/s1 addback^). Parasite pellet and supernatant were separated and 3‘-AMP as substrate added for 1h, P_i_ content was determined with a malachite green based assay. n=4. **B**- Endonuclease activity measured by degradation of double-stranded M13 phage genome on 2×10^6^ total *L. major* statPh promastigotes for 90 min (*L. major*^Cas9/T7/diCre/3‘flox^ *L. major^Δ^*^p1/s1^ clones E10 & F4 and *L. major*^p1/s1 addback^). Pellet and supernatant were separated and dsM13 DNA added for 90 min. DNA degradation was assessed on an 0.8% TAE-agarose gel and semi-quantified using ImageJ. Representative gel shown. Also included are controls containing only parasites and substrate incubated with DNaseI. n=3, Welsh‘s t-test. **C, D**- Endonuclease activity on *L. major* pellet and supernatant fraction was determined as in B but on single-stranded M13 phage genome for the stated reaction time. n=3, Welsh‘s t-test. **E**- Representative gels of C & D.

**Figure 6.**
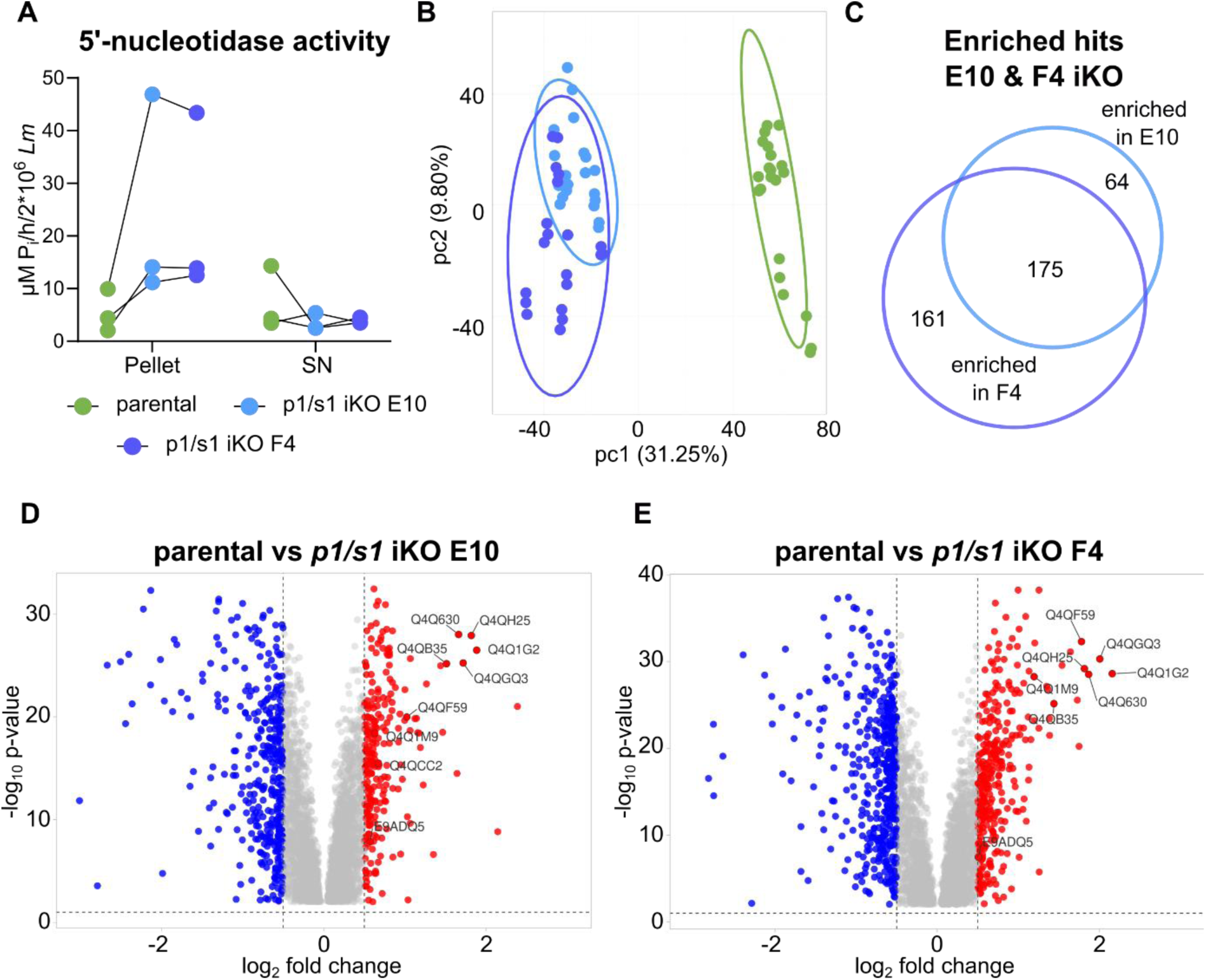
*L. major p1/s1* null mutants bypass loss of p1/s1 via redundant purine salvage. **A**- 5‘-nucleotidase activity was measured on 2×10^6^ intact statPh promastigotes (*L. major*^Cas9/T7/diCre^, *L. major^Δ^*^p1/s1^ clones E10 & F4). n=3. **B**- Principal component analysis of quantitative proteome data generated on *L. major^Δ^*^p1/s1^ clones E10 & F4 and a parental strain (*L. major*^Cas9/T7/diCre/3‘flox^) by label-free mass spectrometry **C**- Venn diagram of enriched proteins in both null mutants. Template created using [43] **D, E**- Vulcano plot of quantified proteins of *L. major^Δ^*^p1/s1^ clones E10 & F4, respectively, in comparison to their parental strain. Cutoffs were set at log_2_FC <-0.5, >0.5 and p-value < 0.01. Denoted are enriched proteins of the purine salvage pathway listed in Tab. 4. **B-E** Data analysis based on n=7 biological replicates in 3 technical repeats each

**Tab. 4.** Purine salvage proteins enriched in both *L. major^Δ^*^p1/s1^ clones compared to parental strain, ranked by p-value in E10 comparison.

| UniProt Code | Annotation | Log<sub>2</sub> ratio E10 | p-value E10 | Log<sub>2</sub> ratio F4 | p-value F4 |
| --- | --- | --- | --- | --- | --- |
| Q4Q630 | Putative 3'-nucleotidase/nuclease | 1.662 | $9.88999 \times 10^{-29}$ | 1.868 | $3.02627 \times 10^{-29}$ |
| Q4QH25 | Nucleobase transporter | 1.819 | $1.24394 \times 10^{-28}$ | 1.816 | $6.61783 \times 10^{-30}$ |
| Q4Q1G2 | Putative membrane-bound acid phosphatase 2 | 1.886 | $3.33012 \times 10^{-27}$ | 2.159 | $2.5267 \times 10^{-29}$ |
| Q4QGQ3 | Putative 3'-nucleotidase/nuclease | 1.719 | $5.82386 \times 10^{-26}$ | 2.006 | $5.09025 \times 10^{-31}$ |
| Q4QB35 | Membrane-bound acid phosphatase 2 | 1.514 | $6.83801 \times 10^{-26}$ | 1.44 | $7.14518 \times 10^{-26}$ |
| Q4QF59;<br>Q4QF58 | Putative nucleoside transporter 1 | 1.023 | 1.05811x10<sup>-20</sup> | 1.78 | 5.1277x10<sup>-33</sup> |
| Q4Q1M9 | Inosine-guanosine transporter | 0.63 | 3.89054x10<sup>-19</sup> | 1.195 | 5.58245x10<sup>-29</sup> |
| E9ADQ5 | Exonuclease domain-containing protein | 0.575 | 1.37787x10<sup>-08</sup> | 0.508 | 1.23532x10<sup>-07</sup> |

### *L. major* can compensate for *p1/s1* loss by enrichment of purine salvage proteins

Essentiality of the *p1/s1* cluster remains elusive, as conventional and facilitated CRISPR-Cas9 methods were lethal to the parasites (n=6) but inducible gene deletion using the same sgRNA templates yielded viable mutants (Tab. 3). This observation raised the hypothesis that the gradual depletion of *p1/s1* during diCre induction allowed adaptation of the parasites to compensate for this deletion. To test this, we measured the parasites’ 5’-nucleotidase activity. This activity, serving as an alternative source of extracellular adenosine for purine salvage, can aid the parasites in supplying metabolite needs. Indeed, we found 5’-nucleotidase activity to be elevated on *L. major^Δ^*^p1/s1^ (Fig. 6A).

To quantitatively assess global proteomic dynamics in *p1/s1* null mutants and expression of further purine salvage-linked proteins, a label-free mass spectrometry analysis was applied. In total, 5890 proteins could be detected, corresponding to 73.28% of the reference proteome.

A principal component analysis revealed similarity between replicates of both *p1/s1* iKO strains, whereas the parental strain clustered separately (Fig. 6B). Successful depletion of p1/s1 was confirmed at the protein level. We found 604 proteins to be differentially expressed between parental strain and iKO E10, 788 between parental strain and iKO F4, with 467 regulated proteins being shared by both comparisons (SuppFig 4A). Focusing only on proteins enriched in *p1/s1* null mutants, 175 are shared by both deletion strains (Fig. 6C). Strikingly, among the proteins strongly enriched in *L. major^Δ^*^p1/s1^ strains are enzymes with redundant 3’-nucleotidase-nuclease activity, membrane-bound phosphatases and purine transporters (Fig. 6D-E; Tab. 4). Gene ontology terms for purine nucleotide binding, adenyl nucleotide binding, and nucleotide binding were significantly enriched *p1/s1* iKO E10 (SuppFig 4B). All of these hits can be involved in purine salvage by nucleic acid and nucleotide catabolism or purine precursor transport, providing the parasites with purines despite their reduced 3’-nucleotidase/nuclease activity. Notably, a xanthine phosphoribosyltransferase (Q4QCC2) as key enzyme in intracellular *Leishmania* purine metabolism [53], was enriched in *p1/s1* iKO E10. As no copy number variations were detected in the genomic loci of these compared to a parental strain (NCBI SRA Accession SUB15491365), upregulation by genomic adaptation is unlikely. This rather points to a posttranscriptional regulation, as typical for *Leishmania*.

### Loss of p1/s1 does not impact cell death hallmarks upon miltefosine treatment

A 3’-nucleotidase/nuclease in *L. infantum* was shown to play a role in parasite susceptibility to miltefosine, an anti-leishmanial drug [54]. As we have identified p1/s1 as enriched in pre-cell death stressed promastigotes, we speculated whether it has a role in parasite cell death, potentially as executer by fragmenting DNA. However, upon treatment with increasing doses of miltefosine, null mutants did not show altered sensitivity compared to the parental or addback strains, neither in viability, phosphatidylserine exposure nor DNA fragmentation (Fig.7A-C). Half-maximal effective concentrations (EC_50_) of miltefosine (Fig. 7D) neither varied for the null mutants nor indicated opposed effect for the addback strain. Thus, p1/s1 does not seem to be involved in intracellular cell death pathways like DNA degradation, and does not play a role in miltefosine sensitivity of *L. major*.

**Figure 7.**
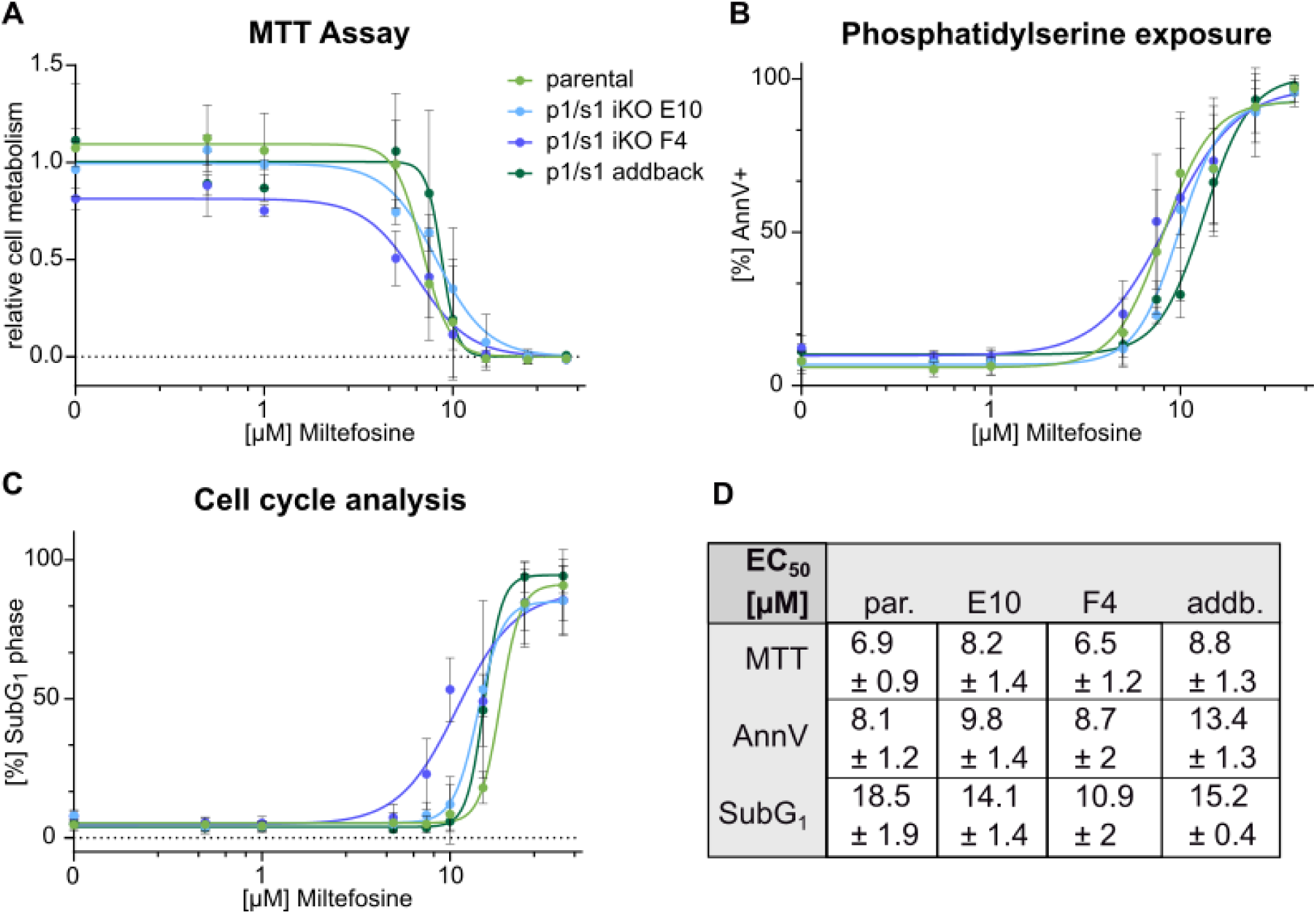
*L. major p1/s1* null mutants do not show altered cell death hallmarks upon miltefosine treatment. *L. major* logPh promastigotes (*L. major*^Cas9/T7/diCre/3‘flox^ *L. major^Δ^*^p1/s1^ clones E10 & F4 and *L. major*^p1/s1 addback^) were treated with increasing concentrations of miltefosine (0.1-60µM) and cell death hallmarks analyzed by different assays. A- Cellular metabolism was quantified using a colourimetric MTT assay, absorbance was normalized to the sample containing 1.5% DMSO as carrier control; n=3. B- Phosphatidylserine exposure on the cell surface was stained using FITC labelled AnnexinV and quantified by flow cytometry; n=3-5. C- Parasites in SubG_1_ phase were quantified by stochiometric DNA staining using SYTOXGreen analyzed by flow cytometry; n=3-5. D- Summarized EC_50_ values obtained from non-linear 4-parameter regression of all cell death assays; ± 95% confidence interval.

### *p1/s1* null mutants cannot decrease PBL proliferation in hMDM:PBL coculture infection and are more susceptible for neutrophil killing

To determine whether p1/s1 serves as virulence factor during macrophage infection, *L. major^Δ^*^p1/s1^ were used in an *in vitro* infection model. In the promastigote stage, *p1/s1* null mutants showed growth deficiency (Fig. 4F) and reduced utilization of extracellular 3’-AMP and nucleic acids (Fig. 5). Presuming that both effects are transferable to intracellular amastigotes, they could lead to reduced parasite persistence in host macrophages. However, in an *in vitro* hMDM:PBL coculture system, infectivity of *L. major^Δ^*^p1/s1^ compared to the parental strain neither decreased in terms of infection rate nor parasite burden (Fig. 8A-B). As shown above, we observed a reduction in PBL proliferation when adenosine or 3’-AMP was added to a hMDM:PBL coculture during infection (Fig. 3C). *p1/s1* deletion alone did not influence absolute PBL proliferation (Fig. 8C). A suppresive effect comparable to Fig. 3C was apparent for the parental strain in the presence of adenosine or 3’-AMP. However, *L. major^Δ^*^p1/s1^ lost the ability to suppress PBL proliferation when 3’-AMP was added (Fig. 8D), likely due to reduced degradation to adenosine. Addition of adenosine as a positive control still reproduced this reduction.

**Figure 8.**
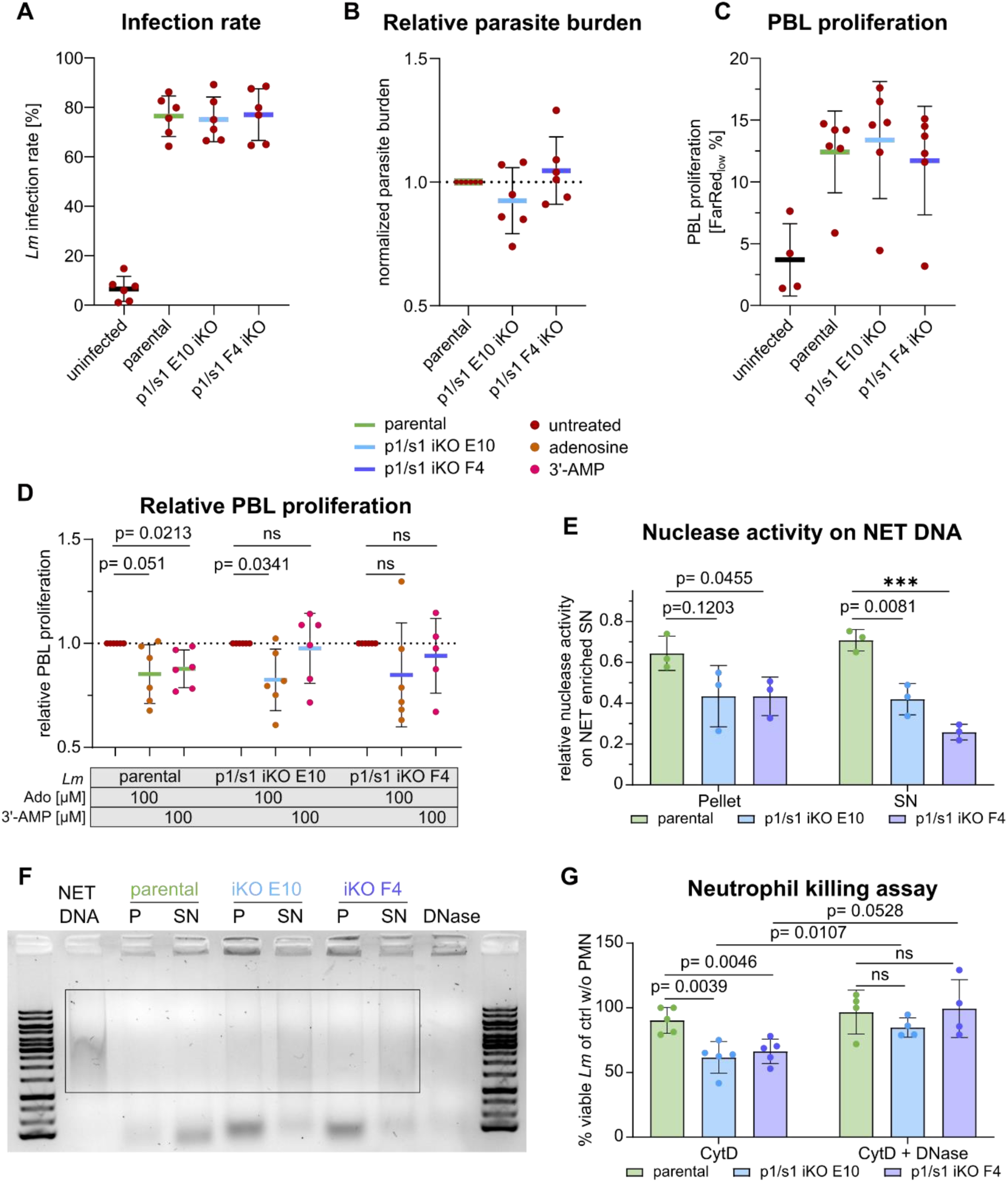
*L. major p1/s1* null mutants cannot decrease PBL proliferation upon presence of 3’-AMP. **A**- *L. major* infection rate of macrophages 4 days pi. Macrophages were infected for 3 hours with MOI=20 of statPh promastigotes (*L. major*^Cas9/T7/diCre/3‘flox^ *L. major^Δ^*^p1/s1^ clones E10 & F4) in the presence of adenosine or 3‘-AMP before autologous peripheral blood lymphocytes (PBLs) were added and incubated for 4 days. Intracellular *L. major* were stained with α-*Lm* mouse serum and labelled with α-mouse-AlexaFluor647 antibody. **B**- Parasite burden measured as MFI of infected macrophages, normalized to the control infected with parental strain. **C**- Percentage of proliferated PBLs was determined as CellTrace_low_ lymphocytes 4 days pi **D**- Normalized PBL proliferation, relative to the respective strain’s infected control without adenosine or 3‘-AMP. **A**-**D**: n=6 donors in 3 independent experiments, Welsh’s t-test **E**- Nuclease activity measured by degradation of human neutrophil extracellular traps (NETs) enriched supernatants on 2×10^6^ total *L. major* statPh promastigotes (*L. major*^Cas9/T7/diCre/3‘flox^ *L. major^Δ^*^p1/s1^ clones E10 & F4). Pellet and supernatant were separated and 500µg NET DNA added for 2 hours. Degradation was assessed on an 0.8% TAE-agarose gel. n=3 donors in three independent experiments. **F**- Representative gel of **E**, box indicates area used for semi-quantification with ImageJ. **G**- *L. major* statPh promastigotes (*L. major*^Cas9/T7/diCre/3‘flox^ *L. major^Δ^*^p1/s1^ clones E10 & F4) were incubated with CytochalasinD treated neutrophils at MOI=1, optionally in the presence of DNaseI. Motile, viable promastigotes were counted for each sample 2 days after infection and normalized to the respective control without neutrophils. n=4-5 donors in 2-3 independent experiments, Welsh‘s t-test.

Besides the p1/s1 3’-nucleotidase activity that potentially counteracts inflammatory signaling by generating extracellular adenosine, the endonuclease activity of p1/s1 can also bring advantages for parasite survival in mammalian hosts. Ecto-nuclease activity of p1/s1 can be beneficial for *Leishmania* promastigotes when encountering neutrophils in the early stages of infection. In response to pathogen detection, neutrophils can extrude neutrophil extracellular traps (NETs) that have antimicrobial capacity. Several pathogens possess extracellular nucleases to evade NET-mediated killing [55–58]. *L. major* was shown to induce NET formation upon neutrophil infection [59, 60]. Whether p1/s1 allows *L. major* to degrade and survive NETs was examined on human primary neutrophils (PMNs) and cell-free NET-enriched supernatants. Control promastigotes of the parental strain were able to efficiently degrade NET DNA, whereas *L. major^Δ^*^p1/s1^ were less able to do so (Fig. 8E-F). This effect was prominent for both parasite pellet and supernatant, comparable to nuclease activity shown in Fig. 5B. To test for antileishmanial effects of NETs on parental and p1/s1 deficient strains, parasites were incubated with PMNs whose phagocytosis was inhibited by Cytochalasin D. The majority of parental strain promastigotes survived this co-incubation but both *L. major^Δ^*^p1/s1^ strains were significantly reduced in number of surviving parasites. When we added DNaseI to degrade NETs, these leishmanicidal effects were lost and survival of the iKO strains resembled that of the parental strain (Fig. 8G).

Taken together, both of the dual p1/s1 activities are able to support evasion of different branches of the innate immune response. Thus, this ecto-enzyme is key in host-*Leishmania* interactions during early infection.

## Discussion

3’-nucleotidase/nuclease activities of several trypanosomatid species have been biochemically characterized and described as ecto-enzymes with manifold functions before [3, 12, 61, 62]. In the present study, we combined quantitative proteomic and genomic editing approaches in *L. major* with human primary *in vitro* infection models to advance knowledge about the importance of p1/s1 in early infection. As enzyme with a dual activity, we were able to link both functions to critical metabolite acquisition as well as to immune escape mechanisms used by the parasite to promote pathogenicity. Furthermore, we utilized quantitative proteomics to dissect how *L. major* adapt to p1/s1 deletion by enriching complementary purine salvage proteins. This highlights the importance of p1/s1 for parasite survival under stress conditions.

Proteomic analysis revealed that p1/s1 is enriched in both drug-induced stress conditions that would eventually lead to cell death at longer incubation. p1/s1 is also more abundant in statPh promastigotes, suggesting that its expression is both stage-specific and inducible. This observation supports the idea that 3’-nucleotidases/nucleases can be induced not only by nutrient depletion [63], but also as part of a stress adaptation or survival mechanism prior to cell death. Possibly, its involvement in nucleotide and nucleic acid catabolism aids the parasites in acquiring essential purine salvage precursors, as 3’-nucleotidase is the dominant ecto-nucleotidase activity in *L. major*.

Recent attention has focused on a 3’-nucleotidase/nuclease as genetic marker for miltefosine resistance in *L. infantum* [54]. However, we did not observe evidence for a direct involvement of p1/s1 in cell death, neither regarding sensitivity of deletion strains towards miltefosine, nor intracellular DNA degradation or phosphatidylserine exposure upon treatment. Thus, a link between miltefosine sensitivity and 3’-nucleotidases/nucleases might be strain specific.

Essentiality of p1/s1 remains unclear since direct or facilitated knockout approaches did not yield viable parasites. In contrast, an inducible deletion method yielded two viable *Δp1/s1* strains, despite the application of identical sgRNA and recombination sequences as in the aforementioned approaches. However, reduced growth of the deletion strains possibly originates from insufficient nutrient acquisition from the culture medium, emphasizing *L. major* dependency on purine uptake. Genome plasticity in *Leishmania* can lead to rapid adaptation in response to *p1/s1* loss during diCre induction, which is not achieved during direct knockout generation [64]. Indeed, comparing quantified proteomes revealed dynamic changes in protein expression. Elevated levels of proteins associated with purine salvage, transport, and redundant enzymes were detected in both *Δp1/s1* strains, similar to the translational adaptation and increased adenosine uptake seen in parasites under purine starvation conditions [65–67]. In fact, we found no evidence for genomic rearrangement in terms of copy number variations of affected genes in the genome of *L. major^Δ^*^p1/s1^ strains, supporting a genome-independent adaptation. These findings can explain how *L. major^Δ^*^p1/s1^ circumvent essentiality of p1/s1 by post-translation compensation strategies to eventually meet their nutrient requirements.

*L. major* lacking the *p1/s1* cluster showed reduced 3’-AMP and nuclease degradation, confirming that both activities directly derive from p1/s1. Differential iKO effects for 3’-nucleotidase and nuclease on the parasite pellet could be linked to *LmjF.30.1460-1490*, predicted to be membrane-bound, possessing only nuclease activity. As expected from literature, substrate preference for ssDNA over dsDNA was apparent, as single-stranded substrate is digested in a much shorter time [3]. Residual enzymatic activity arises from redundant 3’-nucleotidases/nucleases still present in the parasite genome, which are enriched on the protein level.

Besides the predominant expression of the *p1/s1* cluster in the statPh, *L. major* possess three additional putative 3’-nucleotidases/nucleases, *LmjF.12.0400* and *LmjF.31.2300-10*. The two detected in our proteomics data set also tend to be enriched in the statPh. Accordingly, overall 3’-nucleotidase/nuclease activity is higher in statPh than in logPh promastigotes. Enzymatic activity in the supernatant fraction derives from secreted proteins and thus corrobates extracellular localization of p1/s1 [15]. Its secretion is likely caused by an N-terminal signal sequence. These findings align with earlier insights on the *L. donovani* p1/s1 homologue *Ld*Nuc^S^ [8].

Nuclease activity of p1/s1 is not only responsible for degradation of NETs but also allows parasites to escape from them, supporting earlier findings [20]. As published before, NETs do not have leishmanicidal effects on *L. major* parasites, although they are capable of inducing NET formation [58, 59]. However, *L. major^Δ^*^p1/s1^ partially lose the ability to withstand NET-induced killing. Reversion of this effect by addition of DNaseI suggests that this is indeed based on reduced ecto-nuclease activity. Thereby, p1/s1 exhibits NET escape capabilities, as also observed for a *L. infantum* 3’-nucleotidase/nuclease [19].

Ecto-nucleotidase activity of *Leishmania* has a well-established influence on parasite infectivity in mice, suggesting that the generation of extracellular adenosine impacts disease development [37, 68, 69, 70]. This hypothesis is supported by human visceral leishmaniasis patient data, as well as the finding of immunosuppressive effects of adenosine in *Phlebotomus* saliva [71, 72].

In our human primary *in vitro* infection setting, addition of adenosine or 3’-AMP can enhance parasite burden in human macrophages. At the same time, both additives decrease immediate secretion of the strongly pro-inflammatory cytokine TNFα. Previously, we showed that TNFα regulates T cell proliferation in response to *L. major* infection and indeed, addition of adenosine or 3’-AMP to a hMDM-PBL coculture also reduced PBL proliferation [52]. As 3’-AMP is hydrolyzed to adenosine by ecto-3’-nucleotidase activity of *L. major*, these effects directly derive from p1/s1 activity. Accordingly, *Δp1/s1* mutants lose the ability to reduce PBL proliferation in the presence of 3’-AMP.

Taking all findings together, p1/s1 is a multifaceted enzyme involved on several levels during the *L. major* lifecycle. Its dual ecto-activities as 3’-nucleotidase and endonuclease are crucial for nutrient acquisition from hosts, laying the foundations for essential purine salvage. Moreover, p1/s1 is also involved in a vast network of host-parasite interactions, manipulating and shaping innate immune responses during early stages of infection in favor of the parasites. Generation of extracellular adenosine through cleavage of 3’-AMP leads to anti-inflammatory macrophage responses before promastigotes are phagocytosed. Antimicrobial effects of NETs can be circumvented by nuclease activity. However, recent results showed that *L. infantum* with a deletion phenotype for 3’-nucleotidase/nuclease genes have advantages in metacyclogenesis and sand fly transmissibility, marking these enzymes as a double-edged sword [73]. Potentially, the consequences and implications of 3’-nucleotidase/nuclease activity are changing during the *Leishmania* lifecycle, with altered effects in different host environments.

Effects observed in iKO strains are limited due to compensatory capacities of the parasites. Hence, a specific pan-3’-nucleotidase/nuclease inhibitor can have a great potential for further studies on the role of these enzymes for parasite survival and infectivity. Several characteristics of p1/s1 and 3’-nucleotidases/nucleases in general make them interesting as potential drug target: The type I nuclease class is expressed in the infective stages of many protozoa and it has no human equivalent, reducing host toxicity of potential inhibitors [3]. As ecto-enzymes, they are accessible for drug binding and have multiple functions on parasite and host level. Class I nucleases were also shown to be expressed in *Leishmania* amastigotes [8, 14]. Consistently, p1/s1 expression in *L. major* amastigotes is also evident from our proteomics data set. Whether a prompt inhibition is sufficient for *Leishmania* killing or prone to resistance emergence, needs to be studied. Development of a specific inhibitor can be key to determine the role of 3’-nucleotidases/nuclease in established infection as well as studying mechanisms of transmission benefits seen in deletion strains [73]. To accelerate development of such an inhibitor, we are now screening for compounds binding into the predicted active site of p1/s1.

By presenting a comprehensive study on p1/s1, a bifunctional 3’-nucleotidase/nuclease in *L. major*, we elucidated the central role of this ecto-enzyme to emphasize its manifold involvements during infection onset. Our findings highlight the potential of class I nucleases as drug targets, a potential that could be further realized by elucidating the function of p1/s1 in intracellular amastigotes. Overall, 3’-nucleotidases/nucleases in *Leishmania* are highly relevant to study in many aspects along the lifecycle and are key to understand molecular dynamics of *Leishmania* infection.

## Acknowledgments

We thank M-S. Philipp for excellent technical support for flow cytometry experiments and T. Bauer for providing PMNs. Generative AI was only used to enhance comprehensibility and language flow, authors take full responsibility of the content.

## Funding

This research was funded by the LOEWE Center DRUID (projects D3 to G. v. Z. and Core Facility Chemoinformatics & Molecular AI to P. K.), German Research Foundation priority program SPP 2225 project no. 446496937 to G.v.Z. ZA 1176/12-2, project no. 446605368 to U.D. and DFG project no. 318346496 - SFB1292/2 TP11 to U.D.

